# Large-scale non-targeted metabolomic profiling in three human population-based studies

**DOI:** 10.1101/002782

**Authors:** Andrea Ganna, Tove Fall, Samira Salihovic, Woojoo Lee, Corey D Broeckling, Jitender Kumar, Sara Hägg, Markus Stenemo, Patrik K.E. Magnusson, Jessica Prenni, Lars Lind, Yudi Pawitan, Erik Ingelsson

## Abstract

Non-targeted metabolomic profiling is used to simultaneously assess a large part of the metabolome in a biological sample. Here, we describe both the analytical and computational methods used to analyze a large UPLC-Q-TOF MS-based metabolomic profiling effort using plasma and serum samples from participants in three Swedish population-based studies of middle-aged and older human subjects: TwinGene, ULSAM and PIVUS. At present, more than 200 metabolites have been manually annotated in more than 3,600 participants using an in-house library of standards and publically available spectral databases. Data available at the Metabolights repository include individual raw unprocessed data, processed data, basic demographic variables and spectra of annotated metabolites. Additional phenotypical and genetic data is available upon request to cohort steering committees. These studies represent a unique resource to explore and evaluate how metabolic variability across individuals affects human diseases.

## INTRODUCTION

Metabolomic profiling, or metabolomics, can be described as a holistic approach to the study of low-weight molecules (<1,500 Da) called metabolites. These chemical entities, which are the intermediates or end products of metabolism, serve as direct signatures of biochemical activities and play an important role in many common diseases (Floegel et al. 2013; Stegemann et al. 2014; Wang-Sattler et al. 2012; Wang et al. 2011).

A non-targeted metabolomics approach, as opposite to targeted approaches (Dudley et al. 2010), can be used to simultaneously measure as many metabolites as possible from a biological sample. Ultra-performance liquid chromatography (UPLC) and gas chromatography (GC) coupled with mass-spectrometry (MS) have been the preferred technologies to perform non-targeted metabolomics with high sensitivity, allowing the detection of a large number of metabolites (Buscher et al. 2009).

Recent improvements in instrumental technologies and advances in bioinformatics tools have provided the possibility to perform non-targeted metabolomics on large prospective epidemiological studies with thousands of individuals and hundreds of phenotypes measured (Tzoulaki et al. 2014).

However, partially due to the high cost and complexity in data processing, only few epidemiological studies have undergone large-scale metabolomic profiling. Wurtz and colleagues have used nuclear magnetic resonance to profile more than 13,000 individuals and identify four novel biomarkers for incident cardiovascular disease in (Wurtz et al. 2015). Menni and colleagues have integrated metabolomics and epigenetic profiling to evaluate the biological mechanisms of aging (Menni et al. 2013b) and diabetes (Menni et al. 2013a). Our group has integrated genetics and metabolomics data to identified four lipid-related metabolites with evidence for clinical utility, as well as evidence for a causal role in coronary heart disease development (Ganna et al. 2014b).

Here, we present one of the largest UPLC-MS-based metabolomic profiling efforts to date using plasma and serum samples from participants from three Swedish population-based studies: TwinGene, ULSAM and PIVUS, including more than 3,600 participants. Thus far, more than 200 metabolites have been manually annotated using an in-house spectral library of authentic standards and publically available spectral databases. Moreover, thousands of metabolic features, not yet annotated, have been identified across the three studies.

In addition to metabolomic profiling, information on lifestyle, anthropometrics and demographics, and measurements of established cardiovascular biomarkers are available in all participants. In two of the three studies, extensive measurements of subclinical cardiovascular disease are also available. Furthermore, all participants have been linked with national Swedish registries, allowing the identification of incident disease events for a maximum of up to 20 years of follow-up. This resource is used to perform metabolome-wide association studies (MWAS), to explore networks and pathways of metabolites, and to evaluate new methodologies for metabolite annotation. This data was successfully used by our group in a first analysis for association with incident coronary heart disease in 1,028 individuals (131 events) with validation in 1,670 individuals (282 events) (Ganna et al. 2014b).

This article aims to add to the previous publication (Ganna et al. 2014b) by expanding the description of the data acquisition, feature identification and annotation to known metabolites as well as a discussion on the methodological aspects of using metabolomic profiling in large datasets. The included description of the data analysis pipeline can act as a blueprint for researchers in this rapidly increasing field. Furthermore, it contains notes for those that would like to use the data themselves.

## MATERIAL AND METHODS

The methods described in this section are expanded versions of descriptions in previous work (Ganna et al. 2014b).

### Study populations and sample collection

The distribution of baseline characteristics and main cardiovascular risk factors for the individuals included in the three studies are shown in **Table 1**.

**Table 1.** Baseline characteristics of the individuals of the three cohorts included in the study.

| **Cohort** | **PIVUS**<br>n=968 | **ULSAM**<br>n=1,138 | **TwinGene**<br>n=2,139 |
| --- | --- | --- | --- |
| Age (years) | 70 (0.2) | 71 (0.6) | 69 (8.3) |
| Waist circumference (cm) | 91 (11.6) | 95 (9.60) | 94 (11.9) |
| Systolic blood pressure (mmHg) | 150 (22.7) | 147 (18.7) | 143 (20.2) |
| Diastolic blood pressure (mmHg) | 79 (10.2) | 84 (9.4) | 82 (10.6) |
| BMI (kg/m<sup>2</sup>) | 27.1 (4.30) | 26.3 (3.40) | 26.3 (4.00) |
| LDL-C (mmol/L) | 3.37 (0.87) | 3.90 (0.90) | 3.75 (1.01) |
| HDL-C (mmol/L) | 1.50 (0.41) | 1.28 (0.35) | 1.35 (0.40) |
| Triglycerides (mmol/L) | 1.28 (0.60) | 1.46 (0.79) | 1.43 (0.81) |
| Glucose (mmol/L) | 5.34 (1.61) | 5.79 (1.50) | 5.8 (1.37) |
| Current smoker | 99 (10%) | 227 (21%) | 305 (14%) |
| Lipid-lowering medication | 154 (16%) | 100 (9%) | 363 (17%) |
| Blood pressure-lowering medication | 303 (32%) | 394 (35%) | 581 (27%) |
| Female | 486 (50%) | 0 (0%) | 921 (43%) |

Data are mean (standard deviation) for continuous variables and n (%) for categorical variables. BMI, body mass index; LDL-C, low density lipoprotein cholesterol; HDL-C, high density lipoprotein cholesterol

#### TwinGene

The Swedish Twin Registry is a population-based national register including over 194,000 Swedish twins born from 1886 to 2008 (Magnusson et al. 2013). TwinGene is a longitudinal study within the Swedish Twin Registry initiated to examine associations between genetic factors and cardiovascular disease in Swedish twins. Twins born before 1958 who participated in a telephone screening between 1998 and 2002 were re-contacted between April 2004 and December 2008. Health and medication data were collected from self-reported questionnaires, and a blood sampling kit was mailed to the subject who then contacted a local health care center for blood sampling and a health check-up. Contacts were allowed on Monday to Thursday mornings (and not the day before a national holiday), to ensure that the sample would reach the Karolinska Institutet Biobank in Stockholm the following day by overnight mail. The participants were instructed to fast from 8:00 PM the previous night. A total volume of 50 mL of blood was drawn from each individual by venipuncture. Data on cardiovascular health, medication and death dates were collected through linkage with national registers. In total, 12,591 individuals (55% women) participated in the study.

The blood sampling was done as follows: First, a tube containing Ethylenediaminetetraacetic acid (EDTA) was filled and inverted 5 times immediately. These samples were used for DNA extraction. Second, three gel tubes were filled, inverted 5 times immediately, let standing for 30 minutes for coagulation in room temp, and centrifuged for 10-15 minutes at 3800 rpm. Finally, the serum was tapped from the gel tubes to a collection tube and placed in a transport cylinder. Tubes were sent to Karolinska University Laboratory by overnight post (i.e. in room temperature for at maximum 48 hours) where they were frozen at −80° C for up to eight years until metabolomic profiling.

Metabolomics was performed in a subsample of samples from the TwinGene cohort. Specifically, we utilized a case-cohort design by selecting all the incident cases of coronary heart disease (n=282), type 2 diabetes (n=218), ischemic strokes (n=186) and dementia (n=114) up to 31^st^ December 2010 and a sub-cohort of 1,643 individuals (43% women). The subcohort was stratified on median age and sex, and for each of the four strata, we randomly selected a number of participants proportional to the corresponding number of cases. The subcohort was selected to avoid twins, including almost exclusively unrelated individuals. In total, samples from 2,139 individuals underwent metabolomic profiling.

## ULSAM

Men born between 1920 and 1924 in Uppsala, Sweden were invited to participate at age 50 (N=2,841) in this longitudinal cohort study, which was started in 1970 (Ingelsson et al. 2005), and with a participation rate of 81.7% (N=2,322). Individuals have been reinvestigated at the ages of 60, 70, 77, 82 and 88 years. Information collected includes a medical questionnaire, blood pressure and anthropometric measurements, oral glucose tolerance test and 24-hour ambulatory blood pressure. Data on cardiovascular health, medication and death dates were achieved through linkage with national registers. At age 70, EDTA plasma, citrate plasma, serum and whole blood for DNA extraction were collected from fasting participants, kept on ice for a maximum of 4 hours, and stored at −70° C for up to 20 years until analysis. Additional EDTA plasma was collected during the oral glucose tolerance test in ~600 participants. EDTA plasma samples from the age 70 baseline were used for the metabolomic profiling. In total, samples from 1,138 individuals underwent metabolomic profiling.

## PIVUS

PIVUS is a community-based study where all men and women at age 70 living in Uppsala, Sweden in 2001-2005 were invited to participate (Lind et al. 2005). The 1,016 participants (50% women) have been extensively phenotyped including measurements of endothelial function and arterial compliance, cardiac function and structure by ultrasound and magnetic resonance imaging, evaluation of atherosclerosis by ultrasound and magnetic resonance imaging, seven-day food intake records, electrocardiogram (ECG) analysis, cardiovascular autonomic function and body composition by dual-energy X-ray absorptiometry. Cardiovascular events have been validated through review of hospital records. Blood samples were drawn between 8:00 and 10:00 AM after an overnight fast. Samples were rapidly processed in a refrigerated centrifuge and thereafter kept cool until storage in −70°C. Serum was aliquoted and frozen within two hours and EDTA plasma was aliquoted and frozen within one hour. These samples were stored at −70° C for up to 11 years before metabolomic profiling that was performed in serum samples from 968 individuals.

### Sample preparation

Serum or plasma samples were thawed and 100 μL of serum or plasma were transferred to 400 μL methanol in 96-well format to precipitate proteins (**Figure 1a**). This 80% methanol solution was stored at −20°C overnight, and then centrifuged for 30 minutes at 3800g in 4°C to pellet precipitated protein. The supernatant was aliquoted to separate 96-well plates, sealed using a heat-seal foil, and stored at −20°C until analysis. Plate analysis order was completely randomized for each set of injections, and plates were run in batches of two plates, reflecting the autosampler capacity. Within each set of two plates, the run order was again completely randomized to prevent injection order artifacts. Duplicate injections were performed for all samples, with the second set of injections performed upon completion of the first set of injections for all samples. Each set of injections was randomized independently with respect to plate order, and injection order within a plate.

**Figure 1.** Schematic representation of the workflow used to obtain and process metabolomics data. **a**. Data on both MS and MS/MS were acquired from UPLC-MS. **b**. Peaks were detected for each chromatogram. **c**. Peaks were aligned across samples. **d**. Peaks were grouped across samples. Each peak group is called ‘metabolic feature’. **e**. Metabolic features were log-transformed, sample outliers excluded and data were normalized using ANOVA-type normalization, which accounts for factors of unwanted variation. **f**. Metabolites were identified by matching MS/MS reconstructed spectra with the in-house compound library or using publically available databases. *Photo courtesy of Waters Corporation.

## UPLC-QTOFMS data acquisition

Prior to each batch of two plates of samples, instrument maintenance (cone cleaning, mass calibration, and detector gain calibration) was performed, and a quality control (QC) standard mix was injected. Five injections were performed with 1 μL injections of a 20% methanol solution containing 2 μg/mL each of caffeine, terfenadine, sulfadimethoxime, and reserpine. Only the last three of these five injections were used for QC purpose, and evaluated for retention time (+/− 0.05 minutes), signal intensity (<25% relative standard deviation), and mass accuracy (< 3 ppm) of the four compounds. This approach is designed to prevent acquisition of low-quality data, and acquisition of data from experimental samples was not started until the QC standards passed. The QC steps post-acquisition described in detail below were then used to remove poor injections/samples.

One μL of protein-precipitated serum or EDTA plasma was injected on a Waters Acquity UPLC system. Separation was performed using a Waters Acquity UPLC BEH C8 column (1.8 μM, 1.0 x 100 mm), using a gradient from solvent A (95% water, 5% methanol, 0.1% formic acid) to solvent B (95% methanol, 5% water, 0.1% formic acid). Injections were made in 100% A, which was held for 0.1 min, ramped to 40% B in 0.9 minutes, to 70% B over two minutes, and to 100% B over 8 minutes. The mobile phase was held at 100% B for 6 minutes, returned to starting conditions over 0.1 minute, and allowed to re-equilibrate for 5.9 minutes. Flow rate was constant at 140 μL/min for the duration of the run. The column was held at 50°C, while samples were held at 10°C.

Column eluent was infused into a Waters Xevo G2 QTOF MS fitted with an electrospray source. Data was collected in positive ion mode, scanning from 50-1200 m/z at a rate of 5 scans per second. Scans were collected alternatively in MS mode at collision energy of 6 V, and in idMS/MS mode using elevated collision energy (15–30 V). IdMS/MS (also called MS^E^) allows for an unbiased examination of both precursor and fragment ion mass spectra without additional experiments (Broeckling et al. 2013).

Calibration was performed prior to every batch of 96 samples via infusion of sodium formate solution, with mass accuracy within 1 ppm. The capillary voltage was held at 2200V, the source temp at 150°C, and the desolvation temperature at 350°C at a nitrogen desolvation gas flow rate of 800 L/hr. The quadrupole was held at collision energy of 6 volts. Raw data files were converted to .cdf format using Waters DataBridge software for processing (**Figure 1a**).

### Metabolic feature identification

We used the XCMS software(Smith et al. 2006) to perform peak identification, alignment, grouping and filling (**Figures 1b, 1c, 1d**). This software is implemented in R. An example of the code used for the data processing is available at: https://github.com/andgan/metabolomics_pipeline.

#### Peak detection

We performed peak detection in each chromatogram using the *centWave* algorithm (Tautenhahn et al. 2008) implemented in the *xcmsSet* function (**Figure 1b**). We noted that it is important to determine two instrument-dependent parameters: (1) *ppm,* indicating the mass spectrometer accuracy; and (2) *peakwidth,* indicating the chromatographic peak-width range. The former parameter was set as a generous multiple of the mass accuracy of the mass spectrometer (e.g. 25 ppm if the mass accuracy is 2-3 ppm using a multiple of 10), as previously suggested (Tautenhahn et al. 2008). The latter parameter was set by inspecting the peak-width in several chromatograms. To decide the values of remaining parameters in the *xcmsSet* function, we used an approach based on iterative testing of different settings as discussed in the **Technical validation** section. We also evaluated the quality of the algorithm performance by looking at the plot obtained, for one representative chromatogram, with the *findPeaks* function.

The parameters used for peak detection in the three studies were the following:

- TwinGene: *method="centWave", ppm=25, peakwidth=c(2:15), snthresh=8, mzCenterFun="wMean", integrate=2, mzdiff=0.05, prefilter=c(1,5);*
- ULSAM: *method="centWave", ppm=25, peakwidth=c(2:15), snthresh=6, mzCenterFun="wMean", integrate=2, mzdiff=0.05, prefilter=c(1,5);*
- PIVUS: *method="centWave", ppm=25, peakwidth=c(2:15), snthresh=8, mzCenterFun="wMean", integrate=2, mzdiff=0.05, prefilter=c(1,5).*

#### Peak alignment

Chromatographic shifts over time represent a common characteristic of chromatography-coupled mass spectrometry. Without a proper retention time alignment, peaks representing the same compound would not be correctly grouped across different samples because of retention time drift. We used the *retcorr* function to re-align the samples by correcting the retention time shifting. We used the *obiwarp* algorithm (Prince and Marcotte 2006) (implemented in the *retcorr* function), as it is more stable for large number of samples than the *loess* algorithm, which is the default method in XCMS (**Figure 1c**). This algorithm uses the sample with the largest number of peaks as reference for alignment. Visualization of the extent of correction of retention time for each peak across samples can be obtained from the *retcorr* function and can be informative of batch effects or laboratory issues. The parameters used for peak alignment in the three studies were the following:

- TwinGene, ULSAM and PIVUS: *method="obiwarp",plottype="deviation".*

#### Peak grouping

We grouped the aligned peaks across samples using the *group* function. Each group is called ‘metabolic feature’ or simply ‘feature’ (**Figure 1d**). Three main parameters needed to be determined: (1) *bw,* the retention time deviation to be allowed for grouping; (2) *mzwid,* m/z width to determine the peak grouping across samples; (3) *minfrac*, minimum fraction of samples in each group needed to call it as a valid feature. Simulation approaches based on iterative testing of different settings (see the **Technical validation** section) and the plot obtained from the *group* function were used to determine the right values for these three parameters. We kept the *minfrac* parameter relatively low (below 6%) to allow relatively rare and exogenous (e.g. cotinine, a metabolite of nicotine) metabolites to be included among the features brought forward. However, the disadvantage of using a too low *minfrac* parameter is an increased risk of detecting false-positive features or noise. The parameters used for peak grouping in the three studies were the following:

- TwinGene: *bw=2, minfrac=0.03, max=100, mzwid=0.01;*
- ULSAM: *bw=3, minfrac=0.05, max=100, mzwid=0.01;*
- PIVUS: *bw=2, minfrac=0.05, max=100, mzwid=0.01.*

#### Filling missing features

The fact that metabolic features are not detected in all samples in a cohort might be due to a true lack of signals for certain samples (for example, cotinine should only be detectable in smokers) or, most likely, because some peaks are missed by the peak detection algorithm due to the inherent uncertainty when the intensity is close to the signal-to-noise cutoff. To overcome this problem, we used the *fillpeak* function to impute missing intensities for each metabolic feature. The *fillpeak* function uses the raw data, following retention time correction, to fill the missing intensity values. Notice that, if a feature has truly not been detected, the algorithm will assign a value close to the signal-to-noise threshold. Default parameters were used in all three studies.

### Log_2_ transformation, quality control, normalization

Our data acquisition workflow involved the simultaneous collection of low and high collision energy in alternating scans (Broeckling et al. 2013; Plumb et al. 2006). Thus, for every injection two corresponding data files were generated, rendering four files per individual sample. To perform peak identification, alignment, grouping and filling, we jointly processed MS (low collision energy) and MS/MS (high collision energy) chromatograms. This was done to ensure common features names in both datasets. However, in the following steps only the MS data were used, while the MS/MS data were used for metabolite annotation (see further below).

First, feature intensities were transformed to the log_2_ base scale to approximate normal distribution. Second, potential sample outliers were identified by plotting the total intensity for each sample. Samples exhibiting extreme total intensities might indicate sample degradation or technical errors, and such samples were excluded. Third, an ANOVA-type normalization was used to take into account factors of unwanted variation (Kerr and Churchill 2001; Leek et al. 2010) (**Figure 1e**). In the **Technical validation** section, we show that this normalization performed better than other commonly used normalization approaches. The normalization was done by fitting a linear regression for association between each feature intensity and several factors of unwanted variability. We then used the residuals from the regression as new feature intensities. The factors of unwanted variability can be identified by studying the association between several technical variables and the first principal components. In each study, we adjusted for the following factors:

- TwinGene: *retention time correction, analysis date, storage time, unknown cluster effect;*
- ULSAM: *retention time correction, analysis date, sample collection, plate effect;*
- PIVUS: *retention time correction, analysis date, storage time, season effect.*

Finally, feature intensities were averaged between technical duplicates to reduce the inherent instrumental variability. Features with poor correlation between duplicates were excluded. The correlation threshold for feature exclusion is study dependent and was chosen so that the correlation between technical duplicates was significant with P-value < 0.05 after adjusting for multiple testing, using a Bonferroni correction.

In total, we detected 9,755, 10,162 and 7,522 metabolic features in TwinGene, ULSAM and PIVUS, respectively (**Table 2**).

**Table 2.** Summary statistics of the main characteristics of the three studies

|  | **TwinGene** | **ULSAM** | **PIVUS** |
| --- | --- | --- | --- |
| **Number of individuals included in the final dataset** | 2 139 | 1 138 | 968 |
| **No. of total ion chromatograms \*** | 8 126 | 4 776 | 3 846 |
| **Average no. of peaks detected per chromatogram** | 10 295 | 5 989 | 8 205 |
| **No. of metabolic features before exclusion** | 10 996 | 11 028 | 8 185 |
| **No. of metabolic features after exclusion \*\*** | 9 753 | 10 160 | 7 520 |
| **Mean feature correlation between technical replicates \*\*\*** | 0.38 | 0.46 | 0.43 |
| **Mean coefficient of variation (%) between technical replicates** | 3.70% | 5.20% | 2.91% |
| **N. of annotated metabolites** | 202 | 177 | 189 |

\* Including MS, MS/MS and technical replicates
\*\* Exclusion due to low correlation between technical replicates
\*\*\* Spearman correlation coefficient for one metabolic feature between all first and second replicate samples. The reported value is the average across all the metabolics features.

### Metabolite annotation

Tandem mass spectrometry (MS/MS) using precursor ion selection is a well-established tool to elucidate metabolite structure. In traditional non-targeted metabolomics analysis, global MS analysis is followed by targeted MS/MS experiments to confirm the putative identification of significant features. However, this additional step requires additional analytical work, samples, instrumental time and data processing. Recent developments in mass spectrometer technologies have allowed for the acquisition of both MS and MS/MS simultaneously in the same experiment, by alternating low and high collision energy scans. Using this approach, which is a unique feature of Waters systems, ions are fragmented in an indiscriminate manner (i.e. no precursor ion selection) (Plumb et al. 2006). We have previously shown how to use the correlational relationships in the data to assign the correct precursor ion and reconstruct both idMS and idMS/MS spectra for each feature (Broeckling et al. 2013). R code for generating idMS and idMS/MS spectra is provided at https://github.com/andgan/metabolomics_pipeline.

Once both idMS and idMS/MS spectra are reconstructed, we used several approaches for the annotation of metabolic features to metabolites, reflecting the different levels of confidence suggested by the Metabolomics Standard Initiative [MSI] (Sumner et al. 2007) (**Figure 1f**):

- The first approach (MSI level 1) had the highest confidence and was based on matching accurate mass, fragmentation pattern, and retention time with the in-house spectral library of authentic standards collected under the same experimental conditions.
- The second approach (MSI level 2) was based on spectrum and/or m/z similarities, and a reasonable retention time given the chemical properties, and their annotation relied on information available in public databases.
- The third approach (MSI level 3) used a combination of spectral data and accurate mass to assign the metabolite to a chemical class (without knowing the exact origin of the metabolite), and their annotation to a specific class relied on information available in public databases.
- Finally, when all the other approaches to annotation of the metabolite or metabolite class failed, the metabolite was annotated as “unknown” (MSI level 4).

In total, we detected 109 metabolites with the first approach, 102 with the second approach, and could assign the chemical class to 18 metabolites with the third approach. In the Metabolights repository, we reported the names of metabolites with level 1 and level 2 annotation, and the number of annotated metabolites per cohort is summarized in **Table 2**.

## DATA RECORDS

Three data records, one for each study, have been deposited in Metabolights with accession numbers MTBLS93 (TwinGene), MTBLS124 (ULSAM) and MTBLS90 (PIVUS). Each data record contains five sections:

1. Study design description: This section contains general information about the study characteristics and related publications;
2. Protocols: This section contains a detailed description of the protocol used to collect and process the data;
3. Samples: This section reports the anonymized identification code for each participant, as well as main demographic information (e.g. sex and age);
4. Assay: In this section, all the assays (chromatograms) are reported and linked to the anonymized participant identification code. Each participant should have four assays: MS[repl. 1 and 2] and MS/MS[repl. 1 and 2] (here named MSn).We also report factors of unwanted variability (e.g. batch effects) that have been controlled for during normalization. Note that extended information about the assay characteristics can be downloaded in a .txt format.
5. Study files: In this section, all uploaded assays (chromatograms) can be downloaded in .cdf.gz format. The assays follow this filename format: ‘*analysis date*’_‘*study name*’_‘*increasing number01/02*’. MS files filename end with *01 and MS/MS files, collected on higher collision energy, end with *02. The section also contains a file called *m_’study name’_metabolite_profiling_mass_spectrometry.tsv,* which includes all metabolites that have been identified in the corresponding study. The annotated metabolites are linked to different databases to retrieve the chemical formula and additional information, when available. The retention time and m/z values reported correspond to those observed in our data. It also includes the intensities for each metabolite or metabolic feature across the study participants.

Additional information not shown in Metabolights can be obtained by downloading the entire study or the study metadata. This file can then be open with ISAcreator for editing.

## TECHNICAL VALIDATION

### Description of technical variation

In **Table 2**, we report the main summary statistics about the number of chromatograms, peaks and metabolic features detected in each study. We detected less metabolic features in PIVUS, presumably mostly due to the smaller sample size. We further report measures describing the technical variability of our method. These measures were possible to obtain because each sample has been analysed in non-consecutive duplicates. All three studies were comparable in terms of mean feature correlation and mean coefficient of variation across features; the latter varying from 2.9% in PIVUS to 5.2% in ULSAM. We also report the coefficients of variation obtained from a mixture of four known standards analysed between every analytical batch (192 samples/batch). The mix was analyzed in triplicates, and the relative standard deviation (RSD) of the intensity of the M+H signal for each compound was calculated as the SD divided by the mean and expressed as percent. The mean (range) RSDs over 12 random controls were 2.9% (0.5-8.0) for caffeine, 3.9% (0.8-8.4) for sulfadime, 4.0% (0.7-9.7) for reserpine and 4.0% (0.5-10.2) for terfenadine.

### Determination of optimal XCMS parameters

The parameters used in XCMS to detect, align and group peaks can drastically change the number and quality of the identified features. This aspect is often underappreciated in metabolomics studies and only few prior papers have touched upon this topic. Brodsky and colleagues have shown how the parameter tuning affects the inter-replicate correlation and which parameters are likely to have the largest influence. (Brodsky et al. 2010) The authors of XCMS have suggested parameter values for different UPLC/MS instruments; both in a published paper(Patti et al. 2012) and in the online version of XCMS.(Tautenhahn et al. 2012) However, the settings are both instrumental and study-dependent, and need to be fine-tuned independently for each study.

For each study, we randomly selected thirty participants (thus having 120 chromatograms; including replicate 1 and 2, and both MS and MS/MS) and ran several parameter configurations within a reasonable interval around the suggested values. Specifically, we tried 2,161 combinations of different levels for the following parameters: *snthresh, mzdiff, minfrac, mzwid* and *bw*. The parameter configuration that maximizes the feature correlation between technical replicates is likely to be the best choice. In **Figure 2a**, the average feature correlation between technical replicates is reported separately for each parameter suggesting that a configuration with relatively higher *snthresh* and *minfrac,* and lower *mzwid* and *bw* improved the correlation. The *mzdiff* parameter did not influence the correlation. Similar to our observations, Brodsky and colleagues(Brodsky et al. 2010) also reported that an increase in the *minfrac* and decrease in the *mzwid* parameters were associated with higher feature correlation between technical replicates. To maximize the number of detected metabolic features while maintaining high correlation, a *minfrac* parameter (minimum fraction of sample in each group for calling it as a valid feature) slightly lower than the optimal value was employed.

**Figure 2.** **a**. Results of iterative testing of different parameters for detection, alignment, grouping and filling steps in 30 random individuals (120 chromatograms) from PIVUS. We ran all 2,161 possible combinations of values within the reported ranges for five parameters (*sntresh, mzdiff, minfrac, mzwid, bw*). The reported correlations were the averages of the feature correlations between technical replicates for each parameter, across values of the other parameters. The dot in each panel indicates the value that has been used in the final XCMS settings. The last panel indicates the observed correlation for number of features detected across all possible parameter combinations. **b**. In blue: null distribution of the feature correlations between technical replicates obtained by permutation. In red: observed distribution of feature correlations. The observed correlations are almost always higher than those expected under the null. **c**. Feature distributions across samples in all three studies. The red dots represent the average correlations and the dotted bars represent the ranges between the 5^th^ and 95^th^ percentile of the distribution.

### Comparison of different normalization approaches

We determined which normalization approach worked better in our data by comparing the coefficient of variation and feature correlation between technical replicates (**Table 3**). For simplicity, we performed these simulations using PIVUS data since this was the study with the smallest sample size. The ANOVA-type approach clearly outperformed other normalization methods based on single parameter scaling. This can be explained by the ability of the method to adjust for several factors of unwanted variation, such as analysis date or storage time. After applying the best normalization approach, we plotted the correlation between technical replicates for each feature and compared it with what was expected under the null (**Figure 2b**). The null distribution was obtained by permuting each feature 200 times. Almost all features showed a stronger correlation than expected by chance. Nevertheless, some features had low correlation between technical replicates. Those features with a correlation coefficient of <0.3 had generally lower intensity than those with a correlation coefficient ≥0.7 (mean feature intensity 10.7 and 12.0 respectively, P-value_dif_ < 0.0001).

**Table 3.** Comparison of several normalization approaches in PIVUS

| Normalization Method | Mean feature correlation<br>between technical<br>replicates * | Mean coefficient of<br>variation (%) between<br>technical replicates |
| --- | --- | --- |
| **Raw data** | 0.39 | 4.18 |
| **Median** | 0.42 | 4.67 |
| **Median within analysis date \*\*** | 0.42 | 4.67 |
| **Quantile** | 0.40 | 3.79 |
| **ANOVA-type**, adjusting for: |  |  |
| analysis date | 0.43 | 2.97 |
| analysis date, season | 0.43 | 2.89 |
| analysis date, season, storage time | 0.43 | 3.06 |
| analysis date, season, storage time, amount of retention time correction | 0.43 | 2.91 |

\* Spearman correlation coefficient for one metabolic feature between all first and second replicate samples. The reported value is the average across all the metabolics features.
\*\* Median normalization is performed within each specific analysis date.

Finally, we performed a visual inspection of the metabolic feature distributions across samples in all three studies (**Figure 2c**). All three studies were well normalized, although ULSAM had higher variability than the other two studies.

## Targeted MS/MS analysis to determine the quality of the metabolite annotations

After the entire metabolomic profiling workflow and subsequent metabolite annotations were performed, we evaluated the quality and robustness of our untargeted approach by performing a targeted and confirmatory analysis for a random subset of metabolites which we had annotated with level 1 (n=2) or 2 (n=2). We confirmed all four of these metabolites based on matching accurate mass, fragmentation pattern, and retention time. The targeted mass analysis of the selected metabolites (acetylcarnitine, cortisol, arachidonic acid and docosahexaenoic acid) was performed in the individual sample with the highest concentration of that metabolite from the PIVUS study using the multiple reaction monitoring (MRM) mode by monitoring the transitions between the protonated molecular ion ([M+H]^+^) including molecular ion adducts ([M+Na]^+^, [M+H−H_2_O]^+^, [M+K]^+^) and its product ions (fragments). Although limited to a small subset of the annotated metabolites, with a range of retention time from 0.5 minutes (acetylcarnitine) to ~6.3 minutes (arachidonic acid), these results support the high quality of the workflow and annotation. Further, we have previously validated the level 2 annotation of four metabolites associated with coronary heart disease by targeted mass analysis (LysoPC 18:1, LysoPC 18:2 MG 18:2 and SM 28:1).(Ganna et al. 2014b). Nevertheless, we stress that the annotation quality might decrease at the boundaries of the retention time range and we didn’t specifically investigate this scenario.

### Sharing of individual level phenotypic data

All the raw data, including MS and MS/MS spectra, as well as the processed data for the annotated metabolites have been made available on the Metabolights repository. Age, sex and potential factors of batch effect are also accessible in the repository. Phenotypes from the TwinGene, ULSAM and PIVUS studies, other than age and sex, are not shared in Metabolights due to restrictions in the ethical permits. Nevertheless, data can be made available upon request for researchers who meet the criteria for access to the confidential data. Data from the TwinGene study are available from the Swedish Twin Registry steering committee (http://ki.se/en/research/the-swedish-twin-registry-1; contact). Data from the ULSAM study are available from the ULSAM steering committee (http://www2.pubcare.uu.se/ULSAM/res/proposal.htm; contact). Data from the PIVUS study are available from the PIVUS steering committee (http://www.medsci.uu.se/pivus/; contact). Finally, GC-MS analysis of the same plasma and serum samples, as well as LC-MS analysis of urine from the same participants are ongoing and will be shared with the community as soon as the analyses are ready.

## DISCUSSION

This article can act as a blueprint for data acquisition, feature identification and metabolite annotation in large-scale UPLC-MS metabolomics studies.

### Metabolome-wide association studies

Non-targeted metabolomics can be performed in epidemiological studies to identify new biomarkers of disease. Similar to genome-wide association studies (GWAS) in the field of genomics, it is possible to conduct a non-targeted metabolome-wide association study (MWAS), using metabolites as independent variables (instead of genetic variants as in a GWAS) and a disease or other trait of interest as dependent variable. We suggest performing a univariate analysis (e.g. linear regression for continuous outcomes, logistic regression for dichotomous outcomes or Cox regression for time-to-event outcomes) for each feature. These models are typically adjusted for age and sex; but for specific outcomes, additional biological confounders can be included, depending on the research question. Features with low correlation between technical replicates or those observed in only few samples previous to peak filling should be excluded.

Given the large number of statistical tests performed, correction for multiple testing needs to be considered. False discovery rate (FDR) can be used to control the number of false positives among the metabolites declared significant. This strategy works better in situations where a large number of discoveries are expected, which is often the case with metabolomics data, especially when investigating cardiovascular or metabolic traits. Validation in a separate study is highly recommended due to the inherent technical and biological variability of this type of data. Given the differences in age range, blood collection methods and blood partition (serum and plasma); TwinGene, ULSAM and PIVUS provide an excellent opportunity for performing validation studies. Moreover, we have recently introduced a methodology to determine the expected proportion of findings that can be validated in an external study (called rediscovery rate) and the proportion of these findings that are expected to be false positives (Ganna et al. 2014a).

Genetic information can also be integrated to suggest potential biological mechanisms and, importantly, to determine if the metabolite might be causative of the outcome of interest. This strategy, known as Mendelian randomization, (Sheehan et al. 2008) uses genetic variants, called instrumental variables, to disentangle the confounded causal relationships between intermediate phenotypes (e.g. metabolites) and disease. As an illustration of the above procedures, we have recently performed a MWAS of coronary heart disease and identified four new lipid-related metabolites. In this work, further genetic information was integrated to indicate if the associations were likely to be causal (Ganna et al. 2014b).

### Annotation

Both MS and MS/MS chromatograms were collected simultaneously, by alternating low and elevated collision energy scans. This approach facilitated the annotation of metabolites by using correlational relationships across individuals to reconstruct both indiscriminate (id) MS and id MS/MS spectra for each feature (Broeckling et al. 2013). To our knowledge, this approach has never previously been attempted in this large number of samples. Further, our analytical method allowed estimation and control of instrumental variability, since each sample was analyzed in non-consecutive duplicates.

### New methods to match features across studies

Even if the discovery and replication studies have been analysed in the same laboratory and under the same experimental conditions, the metabolic features detected might be different because of study-specific sampling, storage and handling or due to differences in data processing (e.g. the *minfrac* parameter depends on the number of samples as more features are detected when a larger number of samples are jointly processed). In order to determine whether two features represent the same compound, both m/z and retention time need to be matched. The m/z match can be done within a certain confidence interval, depending on the accuracy of the mass spectrometer (e.g. ±0.02 m/z differences). The retention time matching is more challenging and depends on the retention time correction applied during peak alignment. To our knowledge, there is no established strategy to integrate this alignment information to improve the matching of retention time across studies. Further research on this topic is encouraged and can be done using the reported data downloaded from the Metabolights repository.

### Additional uses

A unique characteristic of our samples is the simultaneous collection of MS and MS/MS data. However, few papers have explored how to use this technology to improve metabolite annotation. Our resource represents a unique opportunity to identify new ways to use MS/MS data to improve annotation and analysis.

Given the large sample size, it is possible to use annotated metabolites to perform correlation-based network analysis. This analysis can be further integrated with biological information from for example KEGG or Recon X (Thiele et al. 2013) to confirm correlations and suggest new potential biological pathways. Methods to integrate biological information with observed data correlation structures can also be developed using this data.

### Strength and limitations of our approach

We believe our study have several important strengths and novel aspects. First, most prior articles describing metabolomics methods are technical, presumes a strong chemical background and do not focus on analyses of large human populations. Here, we present a pipeline to process metabolomics data using vocabulary and concepts that are more familiar to researchers in population-based and clinical research. Second, the processing of the data using XCMS software is important and it is often underappreciated in previous reports. We believe that a clear illustration of the XCMS parameters is of great importance. Third, metabolic feature annotation is a key step, which has not been thoroughly described in the literature. Researchers that use metabolomics data from a commercial supplier (which is the common situation for most epidemiological researchers) might not be aware of this aspect since they do not perform the annotation. Here, we provide a clear illustration of the annotation process and its pitfalls. Finally, our data offer insight about the variability of the metabolome in individuals from the general population, providing incremental information to that available in existing metabolomics databases, such as HMDB, Metlin and Massbank.

We also acknowledge several limitations. The analytical workflow that we describe is based on the experience gathered from a specific type of data and platform; however, general considerations are valid for most metabolomics studies in large human populations, and our workflow can be easily implemented on data coming from other UPLC/MS platforms. Moreover, the workflow that we present is directly transferrable to any set of single channel data (e.g. GC/MS) with the caveat that settings should be adjusted to reflect different instruments (e.g. peak width and mass accuracy). We further recognize that the idMS/MS spectra annotation based on data collected already at the first-pass analysis is a unique feature of our platform and it is not available in most UPLC/MS systems, though the feature grouping methods could be applied to low collision energy in-source MS spectra, which can be informative. However, most of the steps of feature annotation described in our paper can be extended to experimentally generated MS/MS spectra, similarly to what has been described by Zhu and colleagues (Zhu et al. 2013). Finally, the differences in collection methods and storage time (in ULSAM up to 20 years) can be a limitation for certain biological questions, as it may decrease the statistical power to find associations due to weaker correlations of metabolites across studies, although it can be argued that it is an advantage as the generalizability increases when studying samples collected under different conditions.

### Conclusion

Non-targeted metabolomics enables investigation of a large number of biological and clinical questions in different areas. Although these methods are gaining popularity, the methodology to analyze and processed metabolomics data has not been well described. Here, we provide a detailed description of the analytical and computational methods used in three Swedish population-based studies. We made available all the raw unprocessed data, processed data, basic demographic variables and spectra of annotated metabolites at the Metabolights repository. In conclusion, this is a unique resource for evaluating the relationship between metabolic variability and human disease.

## SAMPLES, SUBJECTS, AND DATA OUTPUTS

We uploaded this information as ISA-Tab metadata format to the Metabolights repository.

## ACKNOWLEDGEMENTS

We thank Dr. Alexandra Jauhiainen for helpful insights and comments. Further, we want to extend our thanks to all participants of the TwinGene, ULSAM and PIVUS studies for the kind contribution to science. The computations were performed on resources provided by SNIC through Uppsala Multidisciplinary Center for Advanced Computational Science (UPPMAX) under project b2011036.

## ETHICAL STATEMENT

All participants gave informed written consent and the Ethics Committees of Karolinska Institutet or Uppsala University approved the respective study protocol.

## REFERENCES

Brodsky, L., Moussaieff, A., Shahaf, N., Aharoni, A., & Rogachev, I. (2010). Evaluation of peak picking quality in LC-MS metabolomics data. Anal Chem, 82(22), 9177–9187, doi:10.1021/ac101216e.

Broeckling, C. D., Heuberger, A. L., Prince, J. A., Ingelsson, E., & Prenni, J. E. (2013). Assigning precursor-product ion relationships in indiscriminant MS/MS data from non-targeted metabolite profiling studies. Metabolomics, 9(1), 33–43, doi:DOI 10.1007/s11306-012-0426-4.

Buscher, J. M., Czernik, D., Ewald, J. C., Sauer, U., & Zamboni, N. (2009). Cross-platform comparison of methods for quantitative metabolomics of primary metabolism. Anal Chem, 81(6), 2135–2143, doi:10.1021/ac8022857.

Dudley, E., Yousef, M., Wang, Y., & Griffiths, W. J. (2010). Targeted metabolomics and mass spectrometry. Adv Protein Chem Struct Biol, 80, 45–83, doi:10.1016/B978-0-12-381264-3.00002-3.

Floegel, A., Stefan, N., Yu, Z., Muhlenbruch, K., Drogan, D., Joost, H. G., et al. (2013). Identification of serum metabolites associated with risk of type 2 diabetes using a targeted metabolomic approach. Diabetes, 62(2), 639–648, doi:10.2337/db12-0495.

Ganna, A., Lee, D., Ingelsson, E., & Pawitan, Y. (2014a). Rediscovery rate estimation for assessing the validation of significant findings in high-throughput studies. Brief Bioinform, doi:10.1093/bib/bbu033.

Ganna, A., Salihovic, S., Sundstrom, J., Broeckling, C. D., Hedman, A. K., Magnusson, P. K., et al. (2014b). Large-scale metabolomic profiling identifies novel biomarkers for incident coronary heart disease. PLoS Genet, 10(12), e1004801, doi:10.1371/journal.pgen.1004801.

Ingelsson, E., Sundstrom, J., Arnlov, J., Zethelius, B., & Lind, L. (2005). Insulin resistance and risk of congestive heart failure. JAMA, 294(3), 334–341, doi:294/3/334 [pii] 10.1001/jama.294.3.334.

Kerr, M. K., & Churchill, G. A. (2001). Statistical design and the analysis of gene expression microarray data. Genet Res, 77(2), 123–128.

Leek, J. T., Scharpf, R. B., Bravo, H. C., Simcha, D., Langmead, B., Johnson, W. E., et al. (2010). Tackling the widespread and critical impact of batch effects in high-throughput data. Nat Rev Genet, 11(10), 733–739, doi:10.1038/nrg2825.

Lind, L., Fors, N., Hall, J., Marttala, K., & Stenborg, A. (2005). A comparison of three different methods to evaluate endothelium-dependent vasodilation in the elderly: the Prospective Investigation of the Vasculature in Uppsala Seniors (PIVUS) study. Arterioscler Thromb Vasc Biol, 25(11), 2368–2375, doi:10.1161/01.ATV.0000184769.22061.da.

Magnusson, P. K., Almqvist, C., Rahman, I., Ganna, A., Viktorin, A., Walum, H., et al. (2013). The Swedish Twin Registry: establishment of a biobank and other recent developments. Twin Res Hum Genet, 16(1), 317–329, doi:10.1017/thg.2012.104.

Menni, C., Fauman, E., Erte, I., Perry, J. R., Kastenmuller, G., Shin, S. Y., et al. (2013a). Biomarkers for type 2 diabetes and impaired fasting glucose using a non-targeted metabolomics approach. Diabetes, doi:10.2337/db13-0570.

Menni, C., Kastenmuller, G., Petersen, A. K., Bell, J. T., Psatha, M., Tsai, P. C., et al. (2013b). Metabolomic markers reveal novel pathways of ageing and early development in human populations. Int J Epidemiol, 42(4), 1111–1119, doi: 10.1093/ije/dyt094.

Patti, G. J., Tautenhahn, R., & Siuzdak, G. (2012). Meta-analysis of untargeted metabolomic data from multiple profiling experiments. Nat Protoc, 7(3), 508–516, doi:10.1038/nprot.2011.454.

Plumb, R. S., Johnson, K. A., Rainville, P., Smith, B. W., Wilson, I. D., Castro-Perez, J. M., et al. (2006). UPLC/MS(E); a new approach for generating molecular fragment information for biomarker structure elucidation. Rapid Commun Mass Spectrom, 20(13), 1989–1994, doi:10.1002/rcm.2550.

Prince, J. T., & Marcotte, E. M. (2006). Chromatographic alignment of ESI-LC-MS proteomics data sets by ordered bijective interpolated warping. Anal Chem, 78(17), 6140–6152, doi:10.1021/ac0605344.

Sheehan, N. A., Didelez, V., Burton, P. R., & Tobin, M. D. (2008). Mendelian randomisation and causal inference in observational epidemiology. PLoS Med, 5(8), e177, doi:10.1371/journal.pmed.0050177.

Smith, C. A., Want, E. J., O’Maille, G., Abagyan, R., & Siuzdak, G. (2006). XCMS: processing mass spectrometry data for metabolite profiling using nonlinear peak alignment, matching, and identification. Anal Chem, 78(3), 779–787, doi:10.1021/ac051437y.

Stegemann, C., Pechlaner, R., Willeit, P., Langley, S. R., Mangino, M., Mayr, U., et al. (2014). Lipidomics profiling and risk of cardiovascular disease in the prospective population-based Bruneck study. Circulation, 129(18), 1821–1831, doi:10.1161/circulationaha.113.002500.

Sumner, L. W., Amberg, A., Barrett, D., Beale, M. H., Beger, R., Daykin, C. A., et al. (2007). Proposed minimum reporting standards for chemical analysis Chemical Analysis Working Group (CAWG) Metabolomics Standards Initiative (MSI). Metabolomics, 3(3), 211–221, doi:10.1007/s11306-007-0082-2.

Tautenhahn, R., Bottcher, C., & Neumann, S. (2008). Highly sensitive feature detection for high resolution LC/MS. BMC Bioinformatics, 9, 504, doi:10.1186/1471-2105-9-504.

Tautenhahn, R., Patti, G. J., Rinehart, D., & Siuzdak, G. (2012). XCMS Online: a web-based platform to process untargeted metabolomic data. Anal Chem, 84(11), 5035–5039, doi:10.1021/ac300698c.

Thiele, I., Swainston, N., Fleming, R. M., Hoppe, A., Sahoo, S., Aurich, M. K., et al. (2013). A community-driven global reconstruction of human metabolism. Nat Biotechnol, 31(5), 419–425, doi:10.1038/nbt.2488.

Tzoulaki, I., Ebbels, T. M., Valdes, A., Elliott, P., & Ioannidis, J. P. (2014). Design and analysis of metabolomics studies in epidemiologic research: a primer on -omic technologies. Am J Epidemiol, 180(2), 129–139, doi:10.1093/aje/kwu143.

Wang-Sattler, R., Yu, Z., Herder, C., Messias, A. C., Floegel, A., He, Y., et al. (2012). Novel biomarkers for pre-diabetes identified by metabolomics. Mol Syst Biol, 8, 615, doi:10.1038/msb.2012.43.

Wang, T. J., Larson, M. G., Vasan, R. S., Cheng, S., Rhee, E. P., McCabe, E., et al. (2011). Metabolite profiles and the risk of developing diabetes. Nat Med, 17(4), 448–453, doi:10.1038/nm.2307.

Wurtz, P., Havulinna, A. S., Soininen, P., Tynkkynen, T., Prieto-Merino, D., Tillin, T., et al. (2015). Metabolite profiling and cardiovascular event risk: a prospective study of 3 population-based cohorts. Circulation, 131(9), 774–785, doi:10.1161/CIRCULATIONAHA.114.013116.

Zhu, Z. J., Schultz, A. W., Wang, J., Johnson, C. H., Yannone, S. M., Patti, G. J., et al. (2013). Liquid chromatography quadrupole time-of-flight mass spectrometry characterization of metabolites guided by the METLIN database. Nat Protoc, 8(3), 451–460, doi:10.1038/nprot.2013.004.

